# Dynamics of switching at stall reveals non-equilibrium mechanism in the allosteric regulation of the bacterial flagellar switch

**DOI:** 10.1101/2021.04.16.440105

**Authors:** Bin Wang, Yuhui Niu, Rongjing Zhang, Junhua Yuan

## Abstract

Behavior of the bacterial flagellar motor depends sensitively on the external loads it drives. Motor switching, which provides the basis for the run-and-tumble behavior of flagellated bacteria, has been studied for motors under zero to high loads, revealing a non-equilibrium effect that is proportional to the motor torque. However, behavior of the motor switching at stall (with maximum torque) remains unclear. An extrapolation from previous studies would suggest maximum non-equilibrium effect for motor switching at stall. Here, we stalled the motor using optical tweezers and studied the motor switching with a high time resolution of about 2 ms. Surprisingly, our results showed exponentially distributed counterclockwise (CCW) and clockwise (CW) intervals, indicating that motor switching at stall is probably an equilibrium process. Combined with previous experiments at other loads, our result suggested that the non-equilibrium effect in motor switching arises from the asymmetry of the torque generation in the CCW and CW directions. By including this non-equilibrium effect in the general Ising-type conformation spread model of the flagellar switch, we consistently explained the motor switching over the whole range of load conditions. We expect to see similar mechanism of non-equilibrium regulation in other molecular machines.

## Introduction

Flagellated bacteria, such as *E. coli*, swim in liquid by rotating helical flagellar filaments, each driven at its base by a reversible rotary motor (flagellar motor). The switching of the motor rotational direction underlines the run-and-tumble behavior of bacterial swimming. When all motors on a cell rotate in counter-clockwise (CCW), their filaments form a rotating helical bundle that propels the cell to swim smoothly (a run). When one or more motors switch to clockwise (CW) rotation, the cell tumbles to change direction for the next run [1]. Thus, the two main functions of the flagellar motor are torque generation (to drive the rotation of flagellar filament) and directional switching (to induce run-and-tumble behavior) [2].

The behavior of the flagellar motor depends sensitively on the external loads it drives. Torque generation at different loads is characterized by the torque-speed relation [3-5], and the relations are different in the CCW and CW directions [5,6]. In the CCW rotation, motor torque is approximately constant up to a knee speed, after which it falls rapidly to zero, whereas motor torque decreases linearly with speed in the CW rotation, with similar stall torques and zero-torque speeds for both directions [5]. Motor switching dynamics is also load dependent [7-9]. A key characterization of the motor switching dynamics is measurements of the CCW or CW interval distributions [9-14]. Through systematic measurements of the interval distributions for motors under zero to high loads, under different proton motive force (pmf) conditions, and with different numbers of the torque-generating units (stators), a non-equilibrium effect in motor switching was revealed and was found to be dependent on the motor torque [9]. A non-equilibrium model was proposed to explained the measurements, in which the non-equilibrium effect is proportional to the motor torque. At low torque (low loads, low pmf, or low stator number), the non-equilibrium effect is small and the interval distribution is near exponential. At high torque (intermediate to high load, high pmf, and high stator number), the non-equilibrium effect is manifested in the non-exponential shape of the interval distributions [9,12].

The energy input for the non-equilibrium effect ultimately comes from the energy source of the motor: the proton flux through the stators (driven by pmf) [2,15]. This raises an intriguing question regarding the dynamics of motor switching at stall. The proton flux is zero at stall, indicating zero energy input and thus zero non-equilibrium effect, whereas the previous non-equilibrium model would suggest a maximum non-equilibrium effect at stall as the motor torque is maximum. To resolve this contradiction, we sought to characterized the motor switching dynamics at stall. We developed a method to study the motor switching at stall with high temporal resolution (2 ms), by using optical tweezers to stall the motor.

### Observing motor switching at stall

To study the motor switching dynamics without the interfere of chemotaxis signaling, we transformed a mutant *E. coli* K12 strain HCB901 whose chemotaxis network was defective, with the plasmid pBES38 that expresses CheY^13DK106YW^ (a CheY double mutant that is active without phosphorylation) under control of the isopropyl-β-D-thiogalactoside (IPTG)-inducible promoter Ptrc and constitutively expresses sticky filaments [16]. Cells were cultivated with various amounts of IPTG to adjust the level of CW bias, harvested at optical density of 0.45-0.50, sheared to truncate flagella, and then tethered onto the poly-lysine-coated coverslip by shortened sticky filament. Free rotation of cell body was observed for about 30 seconds with bright-flied microscopy, then the laser for the optical tweezers was turned on to clamp the cell body and stop cell rotating (Fig. 1A&B), and the signal from the back-focal-plane interference indicating the *x* position of the cell body was recorded for about 45 seconds with a commercial quadrant photodiode (QPD). A typical trace of motor switching at stall is shown in Fig. 1C and more examples for motor switching at various levels of CW biases are presented in Fig. S1. When motor exerted CCW torque on the cell body, the optical tweezers would exert an opposite force on cell body to balance the motor torque with the signal of QPD showing a positive value, conversely, there would be a negative QPD signal. To minimize the optical damaging of the laser to the motor and maximize the QPD signal, the trap position was chosen to a suitable distance away from the motor (Fig. 1A). Next, the laser was turned off, and free rotation of the cell body was observed for about 30 seconds. Comparing the speed, switching rate and CW bias before and after the stalling period, we found that the photo-damaging effect of the laser trap on the motor was negligible. When the motor is stalled in either CCW or CW, the power spectrum of its *x* signal trace is a Lorentzian *S*(*f*) = *A*/(1 +(*f*/*f*_*c*_)^2^)where *A* is a constant and *f*_*c*_ is the roll-off frequency [17]. This was confirmed by analyzing a period of the trace during which the motor did not switch, as shown in Fig. 1F. The time resolution for the set-up was determined by fitting the power spectrum with the Lorentzian, resulting in a value (1/(2*πf*_c_)) of about 2 ms. The resolution was sufficient for following the process of the motor completing a switch, which typically lasts more than 10 ms [11]. To assure that the steep changes of the position signals originated from motor switching rather than detection noises, we employed the optical tweezers to stall mutant cells with motors only spinning CCW, and found that the position signals were fluctuating with a small amplitude as shown in Fig. S2. Therefore, our setup was suitable to study motor switching at stall.

**Fig. 1.**
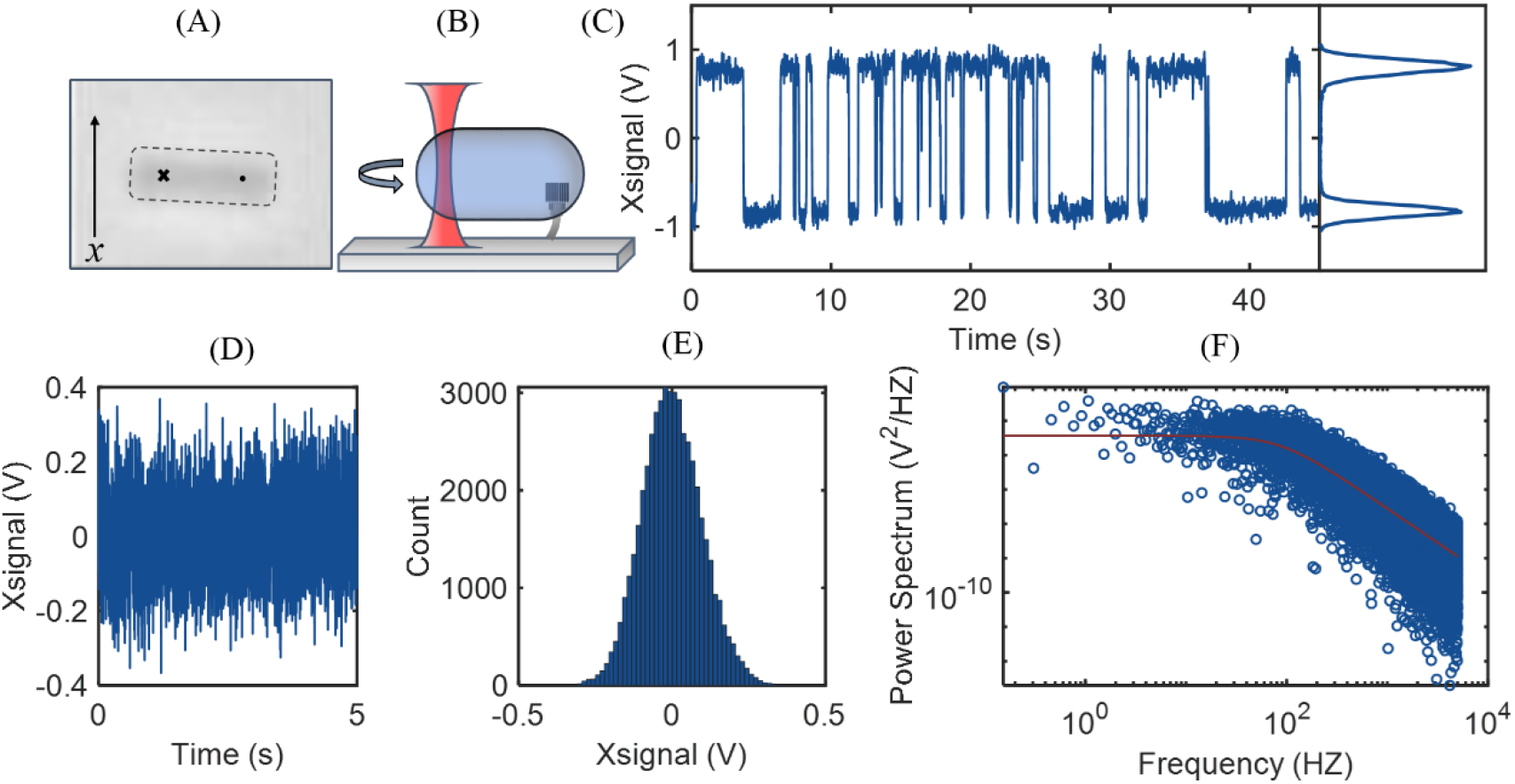
An example of motor switching at stall. (A) The bright-field image of a stalled cell. The dashed line outlines the cell body, the dot denotes the tethered point, and the cross marks the position of the optic tweezers. (B) A schematic diagram of A. (C) A typical trace of the *x*-position signal when the motor switches with CW bias of 0.4. Positive and negative signals indicate CCW and CW torques, respectively. The right panel shows a histogram of the signals. (D) a period of the trace from C when the motor was in CCW state. The averaged signal for the period was subtracted from the signal so that it fluctuates around zero. (E) The histogram of the signal in D. (F) The power spectrum of the trace in D. The red line indicates a fitting with a Lorentzian *S*(*f*) = *A*/(1 +(*f*/*f*_*c*_)^2^) to get the roll-off frequency *f*_*c*_ of 87.6 Hz.

### Interval distributions for motor switching at stall

Using the optical tweezers, 285 motors switching at stall were observed, and the time traces of the QPD signals were converted into binary time series of CW and CCW states, following the procedure described previously [8]. The switching time series were sorted into five groups of CW biases: 0 − 0.2, 0.2 − 0.4, 0.4 − 0.6, 0.6 − 0.8, and 0.8 − 1.0. The interval distributions of CCW and CW states at various CW biases were shown in Fig. 2, exhibiting a near-exponential shape with no apparent peak at short intervals. This is in striking contrast to previous studies where the interval distributions for motors at high torque were non-exponential shape exhibiting a peak at short intervals [9]. With the optical tweezers turned off, the mean motor torques driving the rotation of tethered cells for different groups of CW bias were measured to be 1005 ± 291, 1010 ± 277, 1029 ± 392, 947 ± 330, and 978 ± 293 pN∙nm, which were similar to the high torques in previous experiments [9]. The stall torque, being the maximum torque, is not less than these torques. So we ruled out the possibility that the exponential shape we observed here arose because of low torque. For a positive control, we attached a 750- nm-diameter polystyrene bead to motors of strain HCB901 transformed with plasmid pBES38 and observed with bright-field microscopy. Using the same procedures of sample preparation and data analysis as in the experiments of motor switching at stall, we measured the CW and CCW interval distributions for motor switching under load of 750- nm-diameter polystyrene bead. They are non-exponential shape exhibiting a peak at short intervals (Fig. S3), consistent with pervious experiments at high torque [9]. To make sure the near-exponential shape is not caused by some artifact of optical tweezers, we measured the interval distributions for motors at very high load (under load of 1500-nm-diameter bead). The interval distributions are nearly exponential (Fig. S4), consistent with the measurements at stall. Therefore, motor switching at stall is probably an equilibrium process where the non-equilibrium effect is nearly zero. This is in contradiction to the previous model that proposed that the non-equilibrium effect in motor switching is proportional to the motor torque [9].

**Fig. 2.**
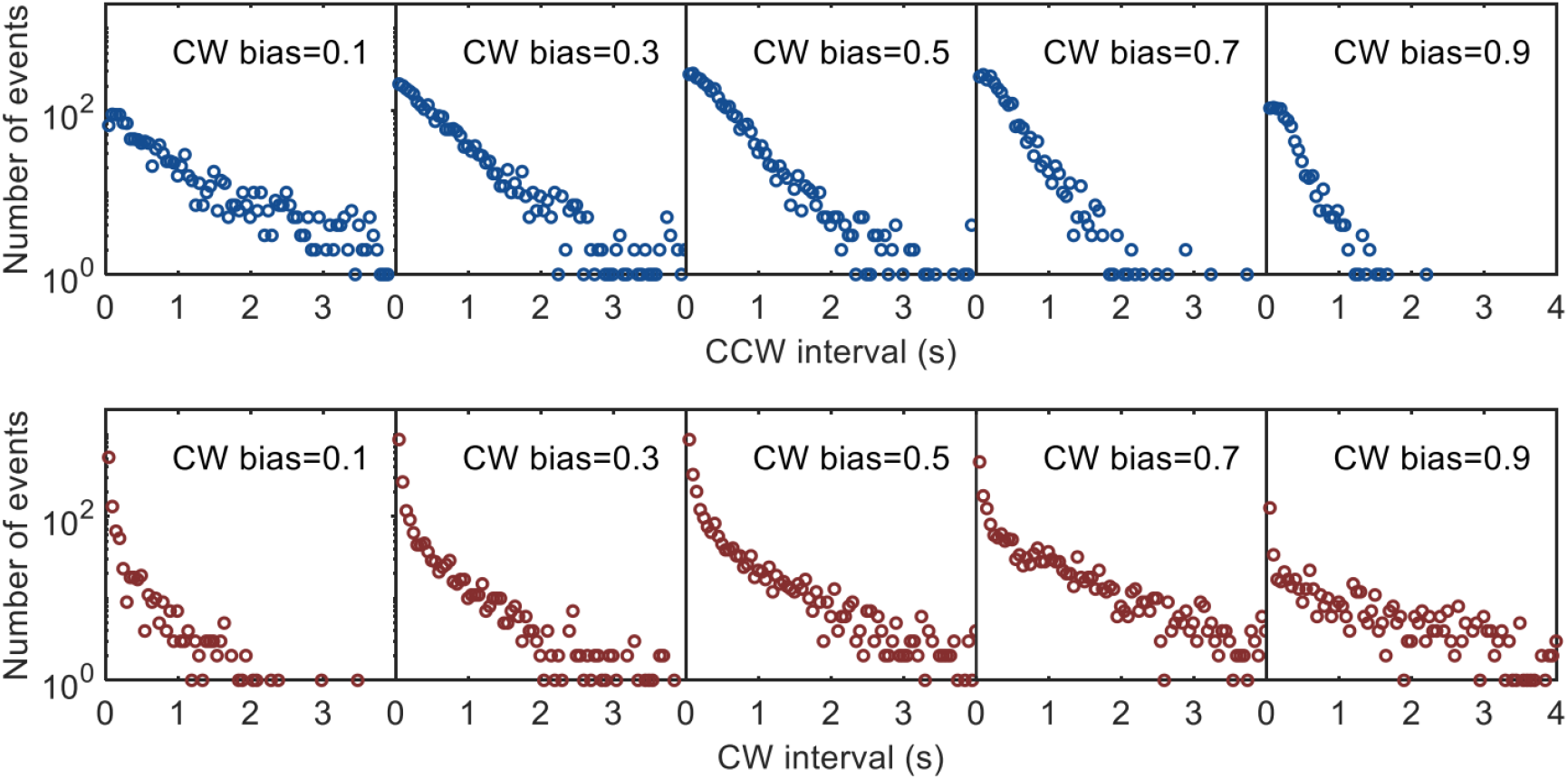
The interval distributions for motor switching at stall. The switching time traces were sorted into five groups of CW biases, and the distributions of CCW (top) and CW (bottom) intervals were plotted. The numbers of motors observed were 39, 55, 71, 76, and 44 for the groups centering on CW biases of 0.1, 0.3, 0.5, 0.7, and 0.9, respectively.

### A new mechanism for non-equilibrium effect in motor switching

To consistently explain the fact that the non-equilibrium effect for motor switching at stall is zero, whereas that for motor switching at other high torques (driving the rotation of micrometer sized beads) is nonzero, a new mechanism for the non-equilibrium effect in motor switching is needed. We sought to introduce the new mechanism for the non-equilibrium effect in a general Ising-type model for the flagellar switch. In the equilibrium conformation spread (Ising-type) model developed by Duke, Le Novere, and Bray [18], a switching event is mediated by conformational changes in a ring of subunits that spread from subunit to subunit via nearest-neighbor interactions. Each subunit exists in four possible states: whether it was activated (CW) or not (CCW), and whether or not it was bound by a CheY-P molecule, as shown in Fig. 3 A. The CCW state was separated from CW state by a free energy difference of *E*_*A*_, and CheY binding would favor the CW state. The free energy changes associated with CheY-binding were *E*_*L*_ − *E*_*A*_ and *E*_*L*_ +*E*_*A*_ for CW and CCW states, respectively. Detailed balance of the transition rates between each pair of states is satisfied in the equilibrium model. To add the effect of motor torque on switching, we expected that the torque would lower the activation barrier between the CCW and CW states by *τδ*, where *τ* is the motor torque and *δ* is an angular constant specifying the torque dependence (Fig. 3B) [8,19]. The transition rates between the CCW and CW states would then increase from their equilibrium values by a factor of exp(*τδ*), where the thermal energy unit *k*_B_*T* was set to be 1. We noticed that the torque-speed relationship is different for CCW and CW rotations as discovered in a recent study (Fig. 3C) [5], with the difference getting larger as the motor speed increase from 0 up to the CCW knee speed of 200 Hz, after which the difference becomes smaller again (Fig. 3D). So there would be a net rate flux from CCW to CW states with the ratio of the two fluxes equaling exp((*τ*_CCW_−*τ*_CW_)*δ*), compared to the equilibrium case where detailed balance was maintained with the ratio equal to 1. Our experimentally measured asymmetry between the shapes of CCW and CW interval distributions for motors at stall further suggested that the angular constants for subunits with (*δ*_B_) and without (*δ*_U_) CheY-P bound are different (details in supporting information), so there will be a net rate flux for each cycle of transitions among the four states, resulting in breaking down of the detailed balance. The extent of broken detailed balance could be seen from the ratio *R* between the products of the transition rate constants along cycles of both directions in Fig. 3E: *R*= *exp*((*τ*_*CCW*_ − *τ*_*CW*_)(*δ*_*B*_ − *δ*_*U*_)). In this new mechanism, the non-equilibrium effect scales with the difference between the CCW and CW torques instead of the average motor torque as in the previous mechanism. This can consistently explain the measurements of motor switching at stall and at other loads. The CCW and CW motor torques at stall are the same, resulting in zero non-equilibrium effect, whereas for motors driving the rotation of micrometer-diameter beads, the difference between the CCW and CW torques leads to non-equilibrium effect in motor switching.

**Fig. 3.**
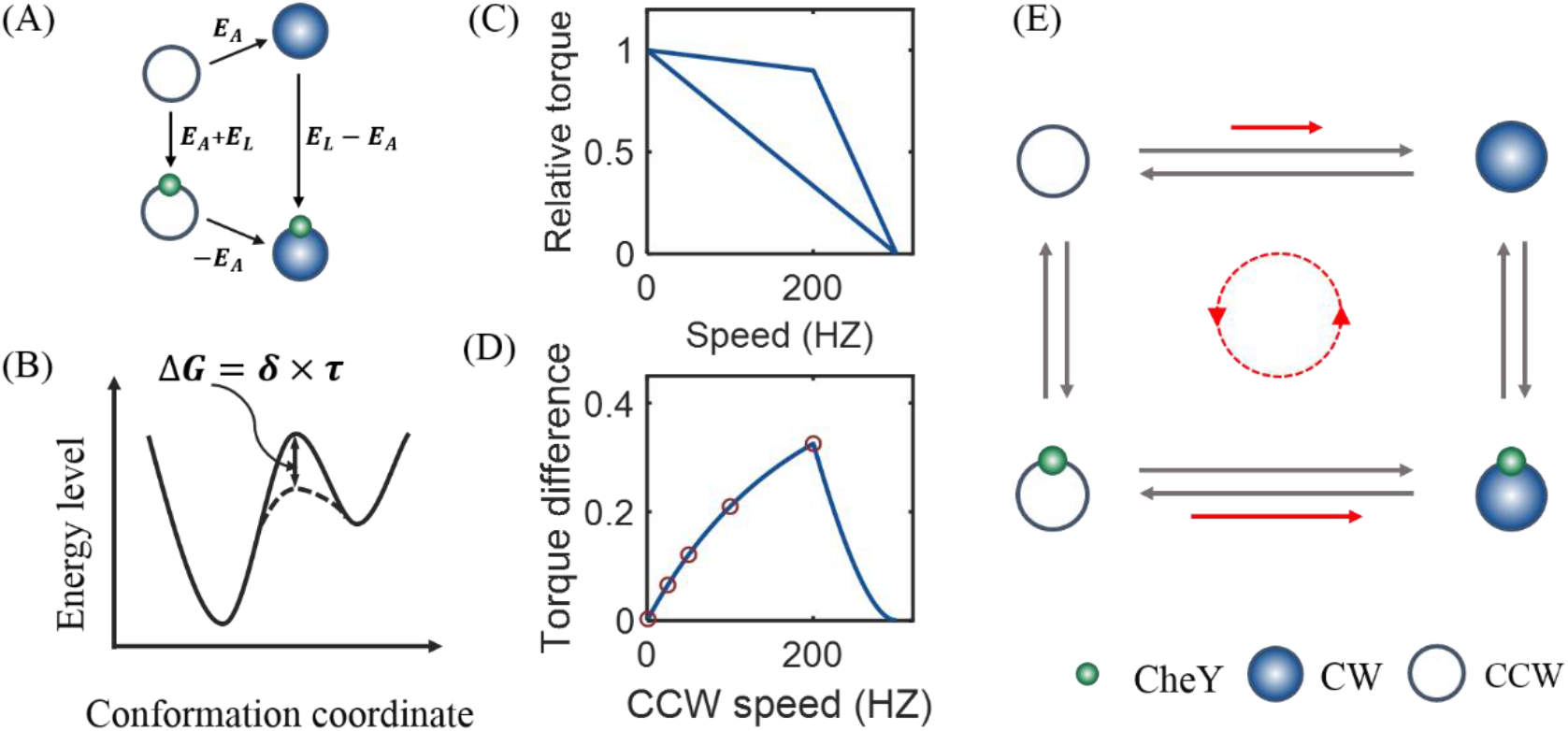
The mechanism of non-equilibrium effect in motor switching. (A) The flagellar switch is modeled as a ring of subunit, each exists in four possible states. The diagram shows the changes of free energy among the states. Open circle, filled blue circle, and small green circle denote CCW state, CW state, and CheY-P molecule, respectively. (B) The stator-rotor interaction lowers the energy barrier between the CCW and CW states by *τ*×*δ* for the subunit that is engaged with a stator, changing the flipping rates between the two states. (C) The torque-speed curves for the CCW (upper curve) and CW (lower line) rotations. (D) The difference in motor torque as a function of CCW speed. The red circles denote the torque differences at speeds of 0, 10, 50, 100, and 200 Hz, which were used in the subsequent simulations. (E) The difference in CCW and CW torques leads to non-equilibrium effect that breaks down detailed balance (red arrows), resulting in a net rate flux (dashed line). The gray arrows indicate the maintenance of detailed balance in an equilibrium model.

We performed stochastic simulation of the non-equilibrium conformation spread model using standard Gillespie algorithm [18,20]. Here for simplicity we consider the symmetrical case where the magnitude of angular constants is the same with or without CheY-P bound but with opposite sign. Using similar values of the parameters as in previous simulations, and with a value for the angular constants (*δ*_B_ = −*δ*_U_) of 0.16, we can reproduce our measured shape of the interval distributions for motors at stall (Fig. 4), and the shape changing from non-exponential to exponential as the motor speed decreases from the CCW knee speed (200 Hz) to stall (Fig. S5). Using *δ*_U_ = −*δ*_B_ = 0.16, the simulated asymmetry in CW and CCW interval distributions at stall was opposite to that experimentally measured (Fig. S6), suggesting that *δ*_B_ > *δ*_U._

**Fig. 4.**
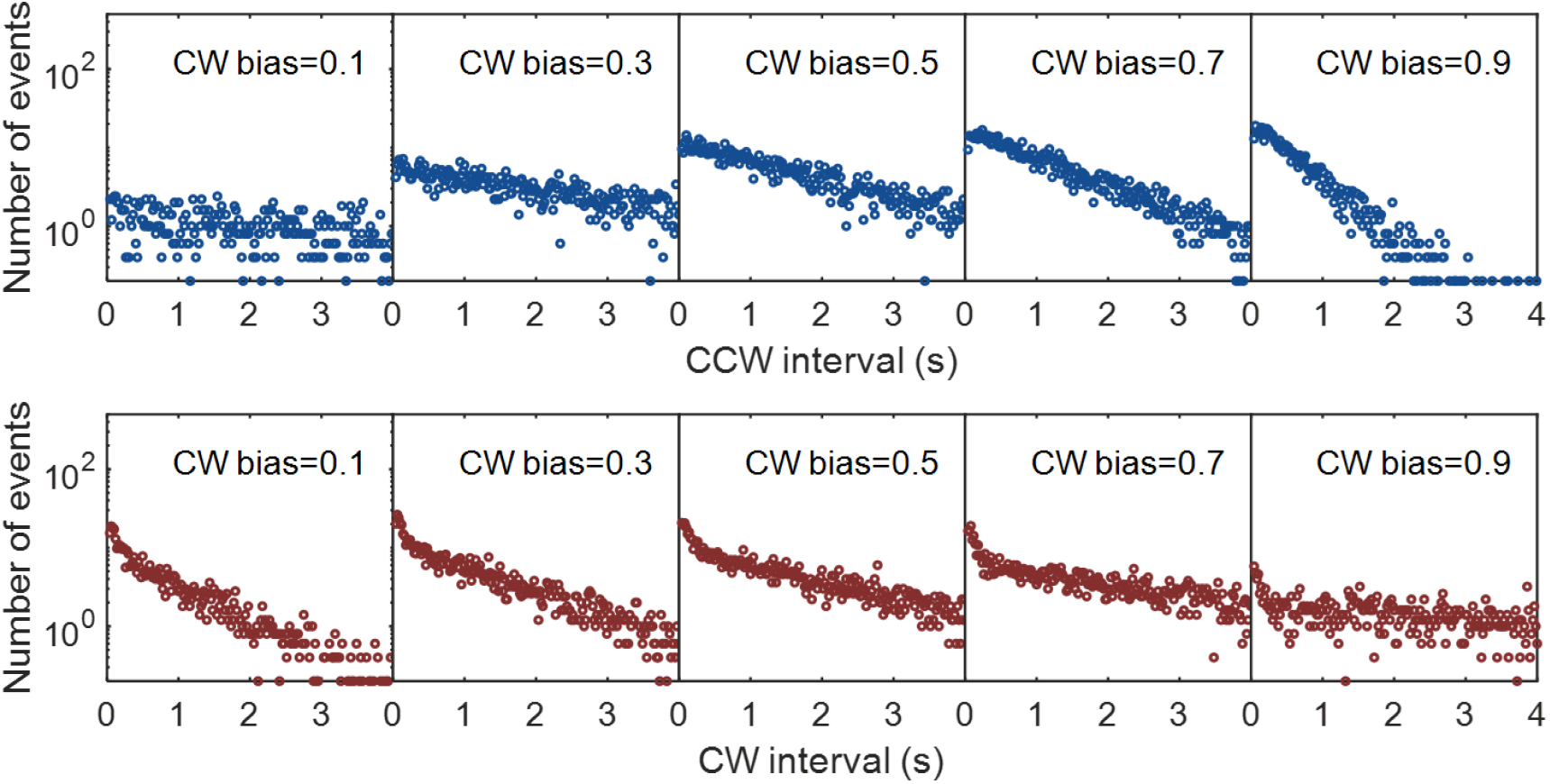
The interval distributions at different CW biases for motor switching at stall from the simulation.

## Discussion

Previous measurements of the stall torque have revealed information about the motor energetics [2,21,22]. Here, we showed that measurement of motor switching at stall revealed further information about the motor dynamics. We developed the method of using optical tweezers to study motor switching at stall with high temporal resolution, finding that the CCW and CW interval distributions for motor switching at stall are exponential shape, which suggested that motor switching at stall is probably an equilibrium process. To resolve the contradiction with the previous model suggesting that the non-equilibrium effect in motor switching is proportional to the motor torque, we identified a new mechanism for the non-equilibrium effect that scales with the difference between the CCW and CW torque. By incorporating the new mechanism into the general Ising-type model of flagellar switch, we consistently explained the experimental results on motor switching dynamics through the whole range of motor torque. In essence, compared with the previous model suggesting that the non-equilibrium effect scales with the motor torque, the new mechanism indicated that the non-equilibrium effect scales with both the motor torque and the relative difference between CCW and CW torques.

The new mechanism started with an assumption similar to previous models [8,19]: the motor torque would alter the energy barrier between CW and CCW states by an amount *τδ* that is proportional to the torque. Detailed balance was maintained with this assumption, thus the models of refs. 8 and 19 are equilibrium models. In our model here, we noticed that the change in energy barrier is *τ*_*CCW*_*δ* when the motor is in CCW state, and *τ*_*CW*_*δ* when in CW state. The extent of broken detailed balance could be seen from the ratio *R* between the products of the transition rate constants along cycles of both directions in Fig. 3E:*R*= *exp*((*τ*_*CCW*_ − *τ*_*CW*_)(*δ*_*B*_ − *δ*_*U*_)). Our measured asymmetry between the shapes of CCW and CW interval distributions for motors at stall further suggested that *δ*_*B*_ > *δ*_*U*_. For motor under all loads except at stall or zero load, *τ*_*CCW*_ > *τ*_*CW*_. Thus *R* > 1 and detailed balance is broken. The magnitude of the scaling constant *δ* corresponds to the angular change in the orientation of the stator-interacting rotor component (FliG) when the motor switches. We could reproduce the experimental results with the model using a value of *δ* of 0.16, which corresponds to an angular change of 9°. Recent cryo-EM maps of the stator and the flagellar switch observed large conformational changes in the stator-interacting FliG component of the switch [23-26]. So this magnitude of angular change is plausible. The finding here suggested an aspect of the functional significance for the asymmetry in torque generation in the CCW and CW rotations [5,6]. The CCW and CW motor torques match at high and low speeds but differ in the intermediate-speed range, with the maximum difference occurring around the CCW knee speed, which is near the load range for a cell swimming in water with the flagellar motor driving the rotation of a full length filament. This results in maximum non-equilibrium energy input to motor switching and lowers the probability of CW/CCW intervals that are too short (< 0.2 s), thereby reducing the occurrence of run durations that are too short and enhancing the efficiency in environmental exploration [27].

We thank Howard C. Berg for strains. This work was supported by National Natural Science Foundation of China Grants (11925406, 21573214, and 11872358), a grant from the Ministry of science and technology of China (2016YFA0500700), and a grant from Collaborative Innovation Program of Hefei Science Center, CAS (2019HSC-CIP004). B.W. is supported by the National Postdoctoral Program for Innovative Talents (BX2018211928).

