## Supporting information for "Dynamics of switching at stall reveals non-equilibrium mechanism in the allosteric regulation of the bacterial flagellar switch"

**The non-equilibrium conformation-spread model**. The model was based on the equilibrium conformation-spread (Ising-type) model developed by Duke, Le Novere, and Bray [18]. The switch complex of the flagellar rotor was modeled as a 1-d Ising ring comprising 34 switching subunits. Each subunit can exist in four possible states as shown in Fig. 3A: two conformational states (CCW/inactive, CW/active), each with two binding states (bound or unbound with CheY-P). The nearest-neighbor interaction was characterized by the coupling energy, which was -$E_{J}$ if the adjacent units were in the same conformational state but zero otherwise. A switching event was mediated by conformational changes in the ring of subunits that spread from subunit to subunit via the nearest-neighbor interactions. For each subunit, the transition rate constant from state *i* to state *j* and the reverse one were proportional to $exp((E_{i}-E_{j})\beta)$, $exp((E_{j}-E_{i})(1-\beta))$, respectively, where the thermal energy unit $k_{B}T$ was set to be 1, and *E_i_* and *E_j_* were the energy levels of state *i* and *j*, $\beta$ was the energy distribution coefficient being always located in [0,1]. Each subunit may bind a molecule of CheY-P with a characteristic binding rate of *k*_L_. The free energy of CheY-P binding to a subunit is *E*_L_= −ln(*c*/*c*_0.5_), where *c* is the intracellular CheY-P concentration, and *c*_0.5_ is the CheY-P concentration at which the motor CW bias is 0.5. *k_A_* was the fundamental flipping rate between CCW and CW states. The changes in free energy associated with CheY-P binding are *E*_L_ + *E*_A_ and *E*_L_ − *E*_A_ for the CCW and CW states, respectively. When a subunit without CheY-P bound is not engaged with a stator, the transition rate from CW to CCW was $k_{A}exp(E_{A}/2)$, and that from CCW to CW was $k_{A}exp({-E}_{A}/2)$. With CheY-P bound, they were $k_{A}exp(-E_{A}/2)$ and $k_{A}exp(E_{A}/2)$, respectively. The CheY-P association rate was $k_{L}\times c/{c_{0.5}}$, and the disassociation rate changed from $k_{L}exp(-E_{A})$ to $k_{L}exp(E_{A})$ when the subunit switched from CW to CCW state. If the subunit was engaged with a stator, the transition rates from CCW to CW and that from CW to CCW were multiplied by a factor of $\exp(\tau_{ccw}\delta)$ and $\exp(\tau_{cw}\delta)$, respectively, where $\tau_{CCW}$ and $\tau_{CW}$ were the motor CCW and CW torque, respectively. The angular constant *δ* for the CheY-P bound and unbound states are *δ_B_* and *δ_U_*, respectively. In the original equilibrium conformation-spread model where the effect of the motor torque is absent, detailed balance was maintained, and the ratio $R$ between the products of the transition rate constants along cycles of both directions in Fig. 3A was equal to 1. In the non-equilibrium model here (Fig. 3E),

$$R=\frac{\exp\left( \tau_{CCW}\delta_{B} \right)}{\exp\left( {\tau_{CW}\delta}_{B} \right)}\times\frac{\exp\left( {\tau_{CW}\delta}_{U} \right)}{\exp\left( {\tau_{CCW}\delta}_{U} \right)}$$

$=\exp\left( {(\tau_{CCW}-\tau_{CW})(\delta}_{B}-\delta_{U} \right))$,

Therefore, the non-equilibrium effect depended on the CCW and CW torque difference, and the difference between the angular constants of $\delta_{B}$ and $\delta_{U}$.

**Evidences for the inequality of** $\boldsymbol{\delta}_{\boldsymbol{U}}$ **and** $\boldsymbol{\delta}_{\boldsymbol{B}}$. For individual C-ring subunits, we calculated the dwelling-time distribution for active (CW) or inactive (CCW) state, leading to $p_{CW}\left( \tau\right)=Aexp\left( -k_{1}\tau\right)+Bexp(-k_{2}\tau)$, and $p_{CCW}\left( \tau\right)=A'exp\left( -k_{1}^{'}\tau\right)+B'exp(-k_{2}^{'}\tau)$, where $k_{1}$,$k_{2}$, $k_{1}^{'}$, and $k_{2}^{'}$ are decay rate constants, and *A*, *B*, *A’*, and *B’* are normalization factors. The formulas for the dwell-time distributions are derived below.

The kinetics of the transitions among the four possible states for each switch subunit is sketched below, along with the rate constants between each pair of states:

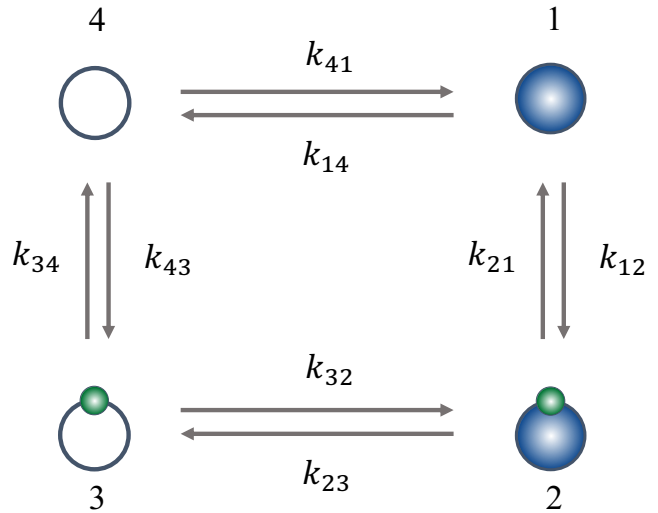

The open circles denote CCW state, the shaded blue circles denote CW state, and the small green circles denote CheY-P molecule. The evolution of the probability vector $p=\left\{ p_{1}, p_{2},p_{3},p_{4} \right\}$ with time was

$\frac{dp}{\mathrm{dt}}=\left( \begin{matrix} -{(k}_{14}+k_{12}) & k_{21} & 0 & k_{41} \\ k_{12} & -(k_{21}+k_{23}) & k_{32} & 0 \\ 0 & k_{23} & -(k_{32}+k_{34}) & k_{43} \\ k_{14} & 0 & k_{34} & -(k_{41}+k_{43}) \end{matrix} \right)p$.

The rate constants are

$k_{12}=k_{L}c/{c_{0.5}}, k_{21}=k_{L}exp(-E_{A})$,

$k_{23}=k_{A}\exp\left( -\frac{E_{A}}{2} \right)\exp\left( \tau_{CW}\delta_{B} \right), k_{32}=k_{A}\exp\left( \frac{E_{A}}{2} \right)\exp\left( \tau_{CCW}\delta_{B} \right)$,

$k_{34}=k_{L}\exp\left( E_{A} \right), k_{43}=k_{L}c/{c_{0.5}}$,

$k_{41}=k_{A}\exp\left( -\frac{E_{A}}{2} \right)\exp\left( \tau_{CCW}\delta_{U} \right), {k_{14}=k}_{A}\exp\left( \frac{E_{A}}{2} \right)\exp\left( \tau_{CW}\delta_{U} \right)$.

Setting the time derivative to 0, we could calculate the steady distribution *p^s^* with the normalization condition of *∑ p^s^ =* 1.

Assume that a subunit switches from CCW to CW state at *t=0,* the survival possibility (*P_cw_*$=\left\{ P_{1}, P_{2} \right\}$) that the subunit is still in CW state at time *t* satisfies

$$\frac{dP_{cw}}{\mathrm{dt}}=M_{cw}{\times P}_{cw},$$

where $M_{cw}=\left( \begin{matrix} -{(k}_{14}+k_{12}) & k_{21} \\ k_{12} & -(k_{21}+k_{23}) \end{matrix} \right)$.

To solve these differential equations, we compute the eigenvalues λ_1_ and λ_2_ of $M_{cw}$. In our model, $k_{A}$ is far greater than $k_{L}$, then $k_{21}k_{12}\ll{(k}_{14}+k_{12}) (k_{21}+k_{23})$, and thus $\lambda_{1}\approx-{(k}_{14}+k_{12})$, $\lambda_{2}\approx-(k_{21}+k_{23})$. Therefore $P_{1}$ and $P_{2}$ would have the bi-exponential form of $C_{1}\exp\left( \lambda_{1}t \right)+C_{2}\exp\left( \lambda_{2}t \right)$， where $C_{1}$ and $C_{2}$ were determined by the initial conditions:$P_{1}^{0}=\frac{p_{4}^{s}k_{41}}{p_{4}^{s}k_{41}+p_{3}^{s}k_{32}}$, $P_{2}^{0}=\frac{p_{3}^{s}k_{32}}{p_{4}^{s}k_{41}+p_{3}^{s}k_{32}}$.

The dwelling-time distribution for the CW state was obtained by

$p_{cw}\left( \tau\right)=-\sum\frac{dP_{cw}}{\mathrm{dt}}=k_{14}P_{1}+k_{23}P_{2}$,

therefore, the shape of the dwell-time distribution for the CW state would be bi-exponential with two decay rates ${k_{1}=-\lambda}_{1}\approx{(k}_{14}+k_{12})$and $k_{2}=-\lambda_{2}\approx(k_{21}+k_{23})$. Using the same method, we would derive the two decay rates $k_{1}^{'}\approx(k_{32}+k_{34})$ and $k_{2}^{'}\approx{(k}_{41}+k_{43})$for the bi-exponential shape of the dwell-time distribution for the CCW state.

At CW bias of 0.5 ($c=c_{0.5})$, and as $k_{L}\ll k_{A},$ we have

$k_{1}\approx k_{A}\exp\left( \frac{E_{A}}{2} \right)\exp\left( \tau_{CW}\delta_{U} \right)$,

$k_{2}\approx k_{A}\exp\left( -\frac{E_{A}}{2} \right)\exp\left( \tau_{CW}\delta_{B} \right)$,

$k_{1}^{'}\approx k_{A}\exp\left( \frac{E_{A}}{2} \right)\exp\left( \tau_{CCW}\delta_{B} \right)$,

$k_{2}^{'}\approx k_{A}\exp\left( -\frac{E_{A}}{2} \right)\exp\left( \tau_{CCW}\delta_{U} \right)$.

At stall,$\tau_{CW}=\tau_{CCW}$. If $\delta_{U}=\delta_{B}$, then $k_{1}=k_{1}^{'}$ and $k_{2}=k_{2}^{'}$. Solving for the normalization factors *A*, *B*, *A’*, and *B’* under the current condition would lead to *A* = *A’* and *B* = *B’*. Therefore, the CW and CCW interval distributions at CW bias of 0.5 for each switch subunit at stall would be the same, suggesting that the motor CW and CCW interval distributions at CW bias of 0.5 would be the same at stall. The experimentally observed asymmetry between the motor CW and CCW intervals at CW bias of 0.5 suggested that $\delta_{U}\neq\delta_{B}$.

**Materials and Methods**

**Strains and plasmids**. All the strains for this study are derivatives of *E. coli* K12 strain RP437. To perform the experiments for motor switching, HCB901(*ΔcheZ fliC*, Ptrc420 ${cheY}^{13DK106YW}$) was transformed with a plasmid pBES38, which constitutively expresses both $\mathrm{LacI}^{q}$and the sticky filament $\mathrm{FliC}^{\mathrm{st}}$. For studying the stability of the setup, strain JY9 (*ΔcheY fliC*, *flgE* modified with CCXXCC at codon 220) with motors spinning only in CCW was transformed with the plasmid pKAF131, which constitutively expresses the sticky filament.

**The procedure to observe motor switching at stall.** Cells of HCB901 with pBES38 were grown at 33 °C with the antibiotics ampicillin (100 μg ml^-1^) and various amounts of inducer IPTG. Cells of JY9 with pKAF131 were also grown at 33 °C with the antibiotics ampicillin at the concentration of 25 μg ml^-1^. Both cells were grown to an OD_600_ of 0.45-0.5, harvested and washed twice with motility medium (10 mM potassium phosphate, 0.1 mM ethylenediaminetetraacetic acid (EDTA), 10 mM lactic acid, and 70 mM NaCl at pH 7.0), and were sheared to truncate flagella by passing washed-cell suspension 200 times between two syringes equipped with 23-gauge needles and connected by a 7-cm length of polyethylene tubing (0.58 mm i.d., No. 427411; Becton Dickinson). Cells were tethered by the shortened sticky filament onto a ploy-lysine coated glass coverslip, which was used for constructing a chamber with a glass slide as the base and two double-sticky tap as the spacer. Untethered cells were washed out by rinsing the chamber with 100 μl of motility medium. The optic tweezers were constructed as described previously [1], by focusing an expanded parallel light beam from a 1,064-nm fiber laser (AFL-1064-33-B-FA; Amonics) into a diffraction-limited spot with a water-immersion objective (Nikon Plan Apo vc 60×/1.20 WI). The optical tweezers were operated to stall the rotation of the tether cell, and the signal from the back-focal-plane interference was recorded by a QPD (InGaAs quadrant photodiodes, G6849, Hamamatsu) for 45 s at a sampling frequency of 10000 Hz. The resulting time resolution for the set-up was about 2 ms. The time resolution could be further improved by using a higher laser power, but to avoid possible photo damage, 150 mW of laser power was chosen in the experiments. Before and after the stalling period, the rotations of the tethered cell were observed with bright-field microscopy with a CMOS camera (Thorlabs, DCC1545M) at a sampling frequency of 100 Hz, each for 30 s. The signal traces were converted to binary time series, using the threshold-crossing algorithm described previously [2]. To measure motor switching under the load of 750-nm-diameter polystyrene bead, the bead was labelled to shortened flagellar stub by passive adsorption, and the rotation of the bead was observed for 200 s with a CMOS camera (Hamamatsu, C11440-22CU) at a sampling frequency of 1000 Hz. The motor torque in tethered cells was calculated using the relation $T_{motor}=\left( \varepsilon_{r}+\varepsilon_{t}{\times s}^{2} \right)\times\omega$, where $\varepsilon_{r}=(8\pi\eta a^{3}/3)/(ln 2a/b-1/2)$ was the rotational drag coefficient of cell body about the short body axis, $\varepsilon_{t}=8\pi\eta a/(ln 2a/b+1/2)$ was the transitional drag coefficient of cell body along the direction perpendicular to the long body axis (the cell body is modelled as an ellipsoid with semi-major axis *a* and semi-minor axis *b*), $s$ was the displacement between the center of cell body and the motor along the long body axis, and $\omega$ was the rotational speed [3].

**Simulation with the non-equilibrium conformation spread model.** The rotor is modelled as a ring of subunits, each existing in either an inactive (CCW) or an active (CW) conformational state with a fundamental flipping rate of $k_{A}$ when no stator is engaged with the subunit. The coupling energy between adjacent subunits is −*E*_J_ if they are in the same states but zero otherwise. The rotor rotates stochastically in a stepwise fashion with a step size of $\Delta\theta=\frac{2\pi}{26}/N$ and a stepping rate of $\frac{\omega}{\Delta\theta}$, where *N* was the stator number (*N* = 10 in our simulation) and *ω* is the motor speed [4]. At a specific motor speed, the CCW and CW torques per stator were determined according to the torque-speed curves in Fig. 3C, with the values of the stall torque per stator, zero-torque speed, and CCW knee speed equal to 200 $pN\cdot nm$, 300 Hz, and 200 Hz, respectively [5]. Ten stators were distributed in average-distributed fixed positions along the ring of rotor subunits. The formulas for the transition rates among different states were presented in the first section of supporting information. The Monte-Carlo simulations were performed with a standard Gillespie algorithm [6]. The values of the parameters used in the simulations are: *E*_A_ = 1.0 , *E*_J_ = 4.0, *k*_L_= 75 s^-1^, *k*_A_ = 10000 s^-1^, *δ*_B_ = −*δ*_U_ = 0.16 (unless otherwise stated).

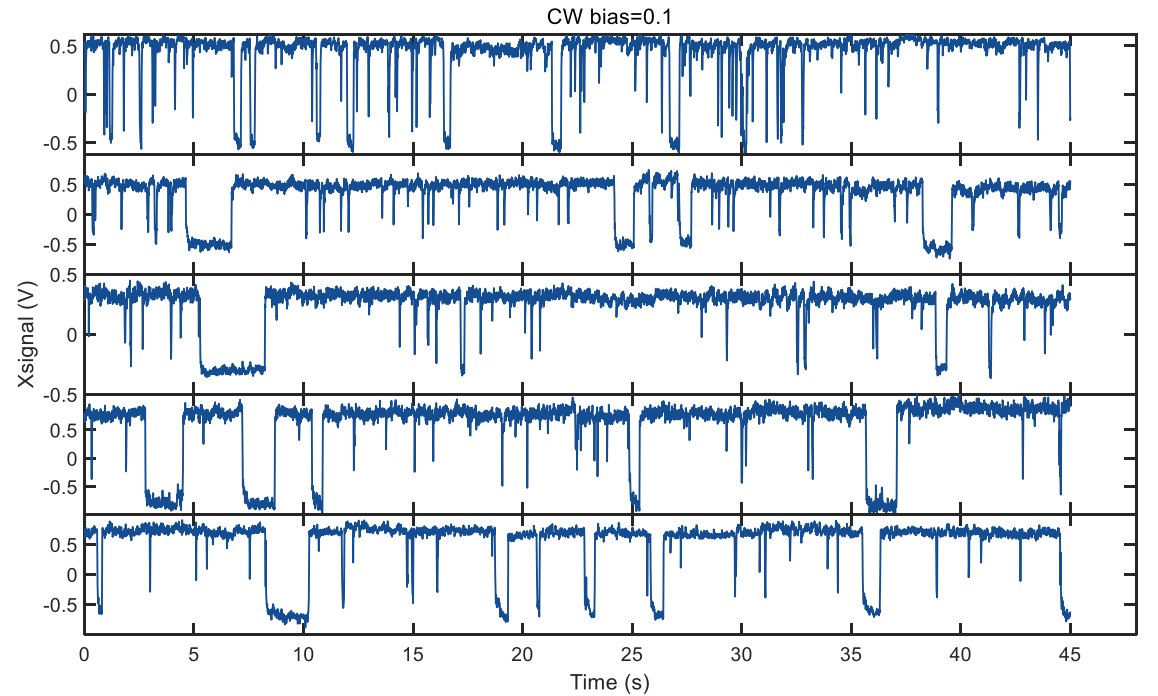

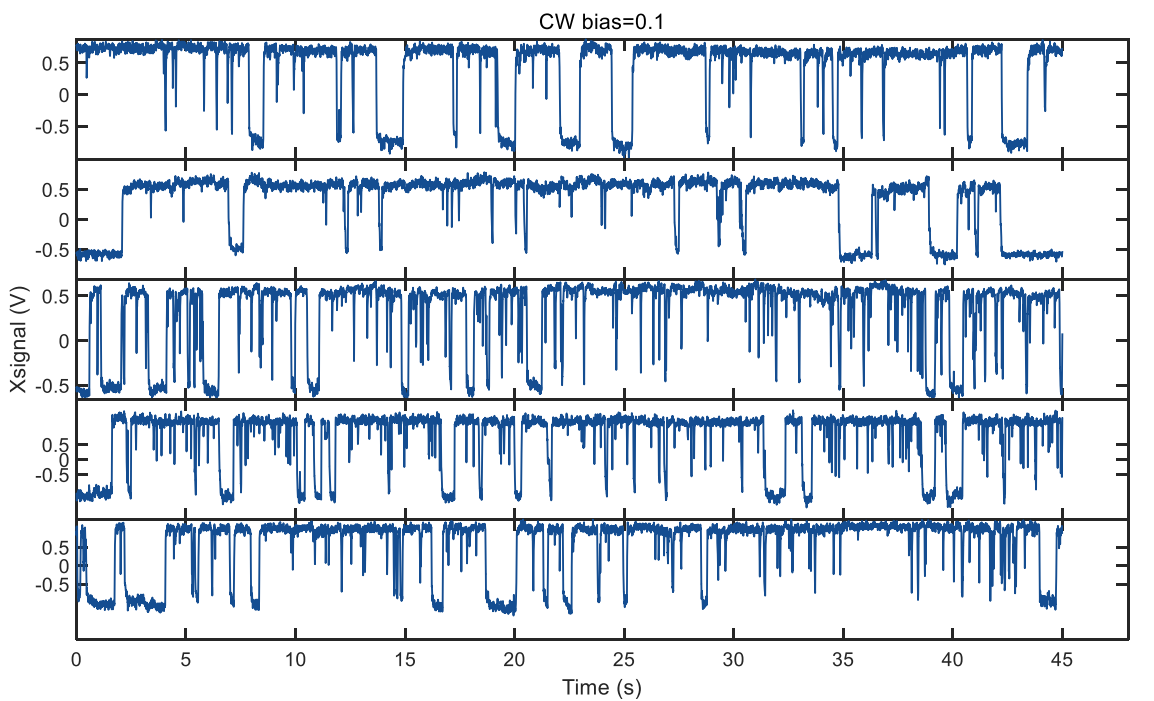

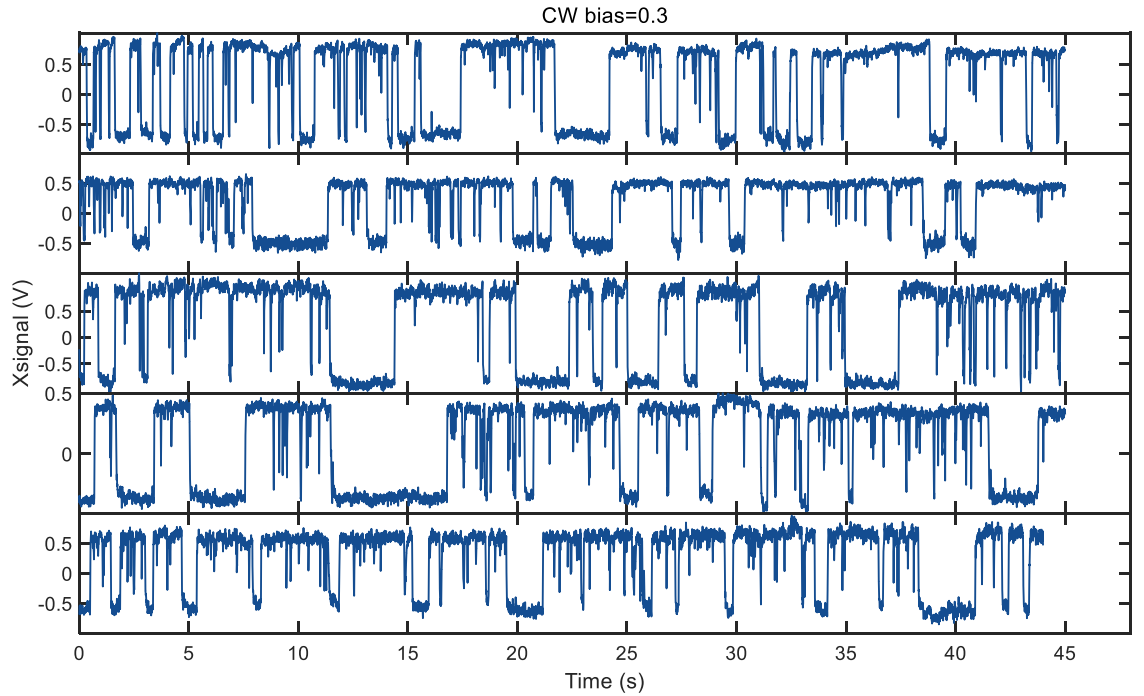

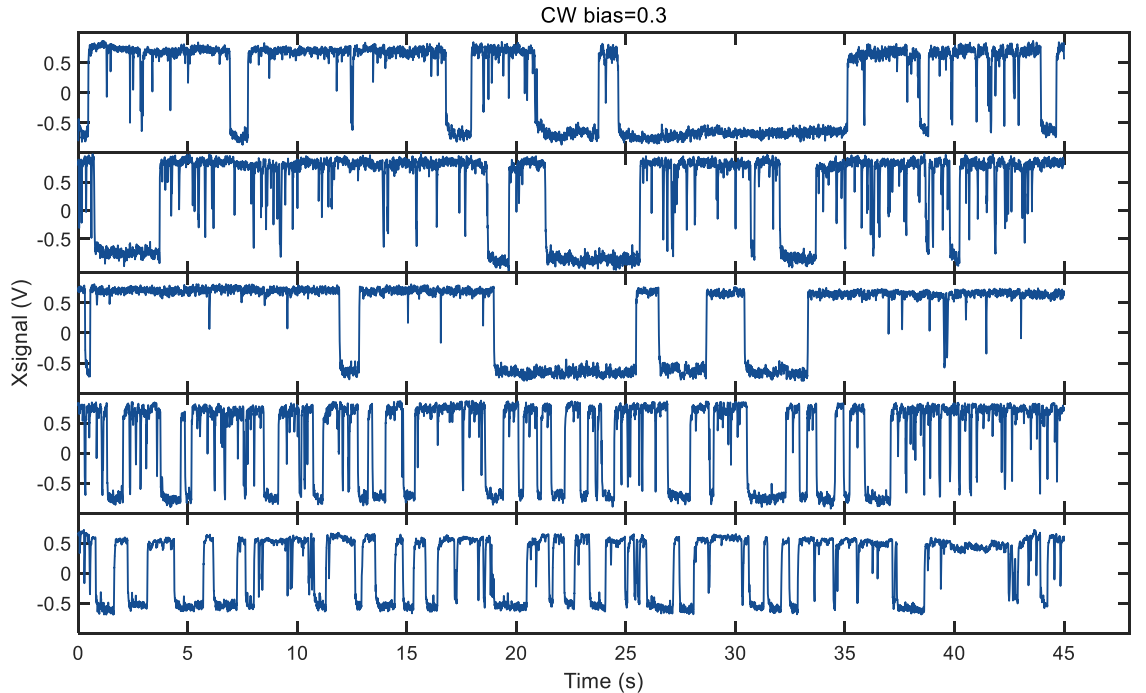

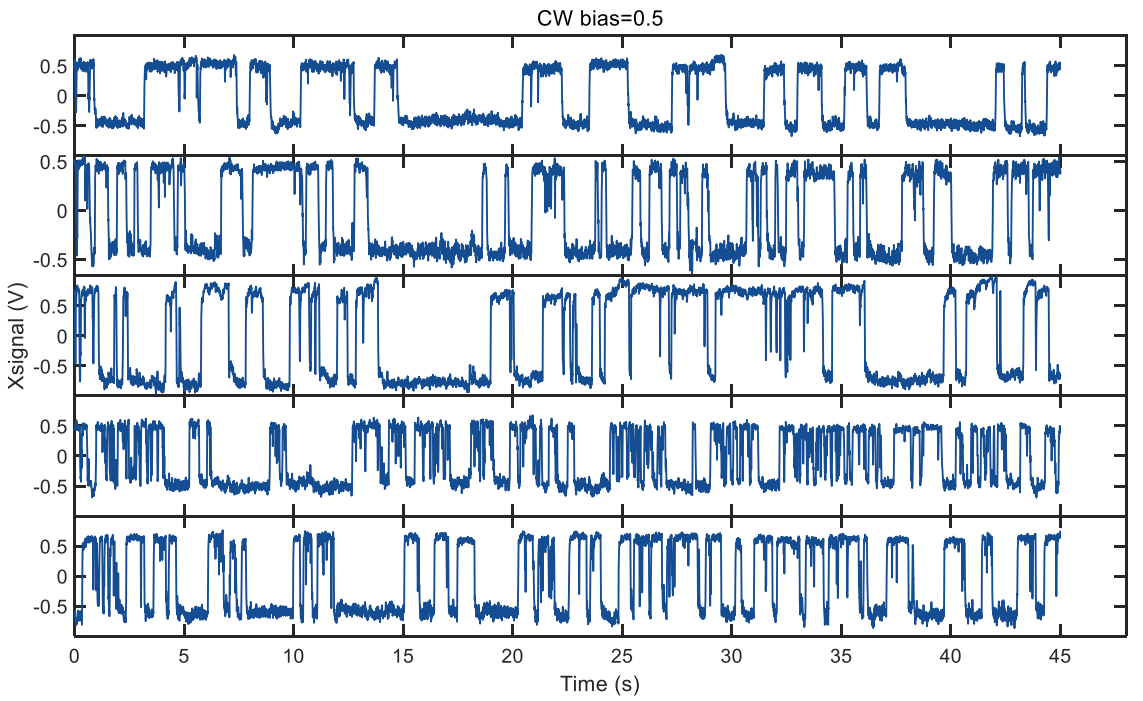

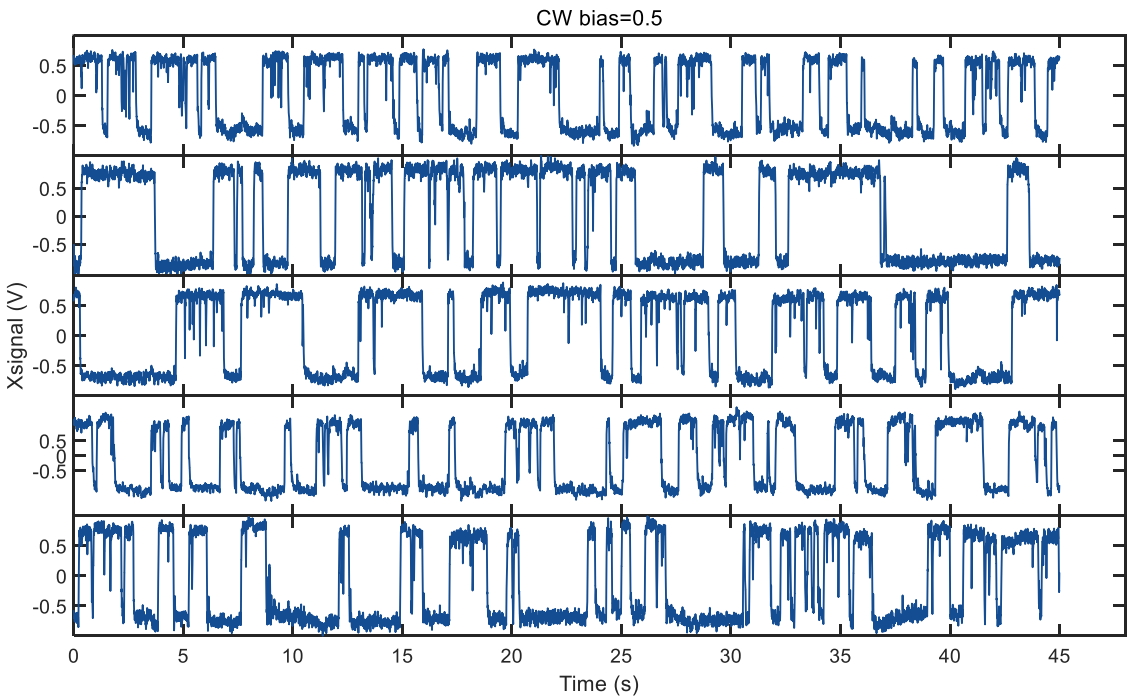

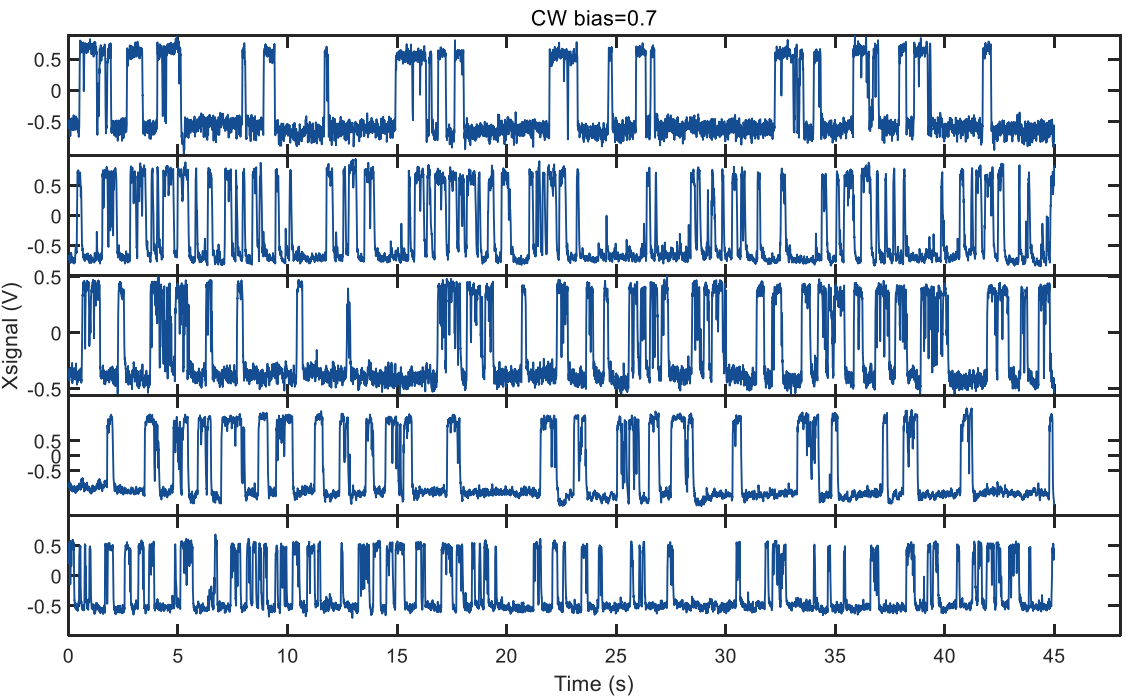

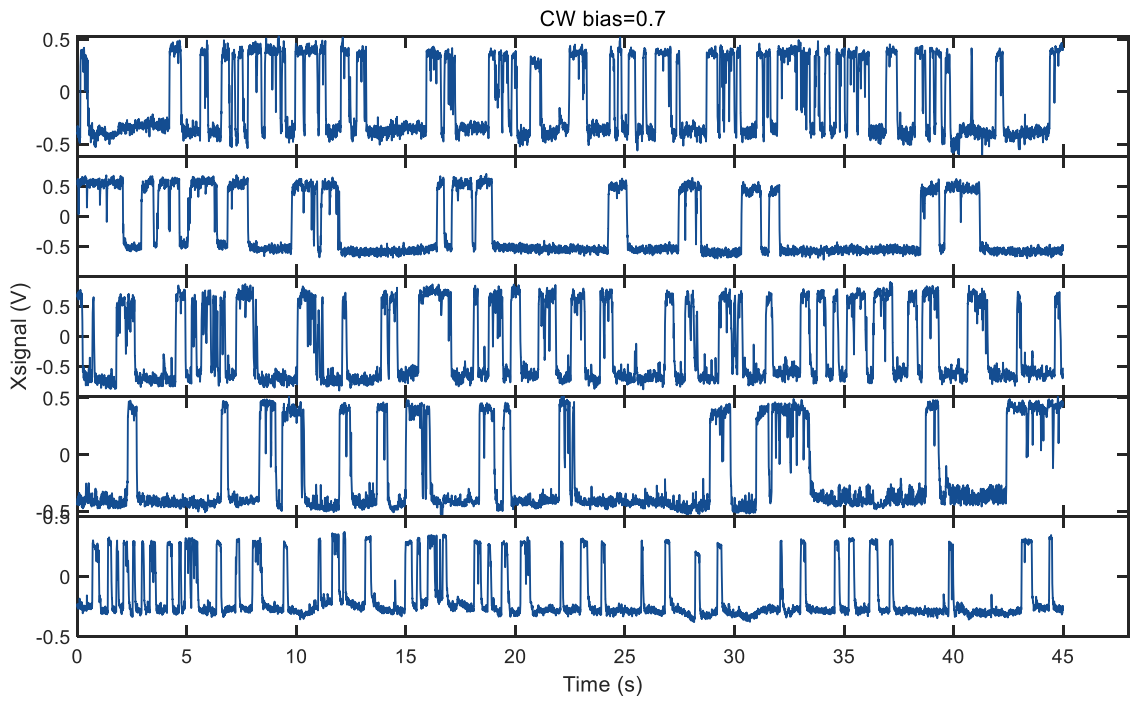

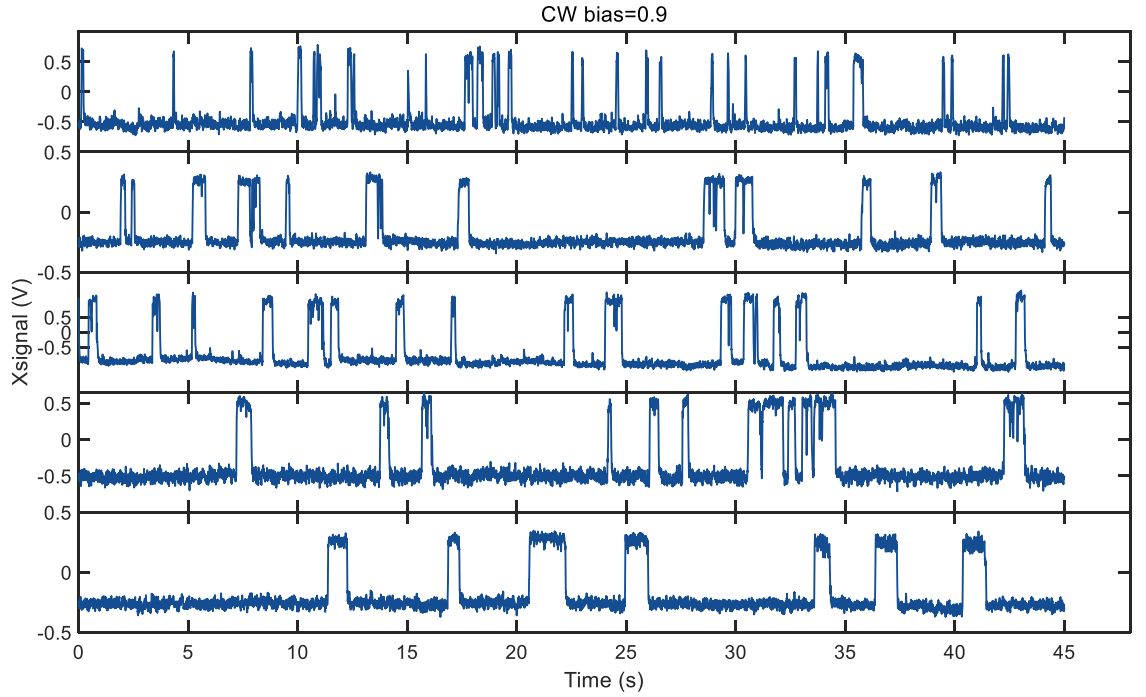

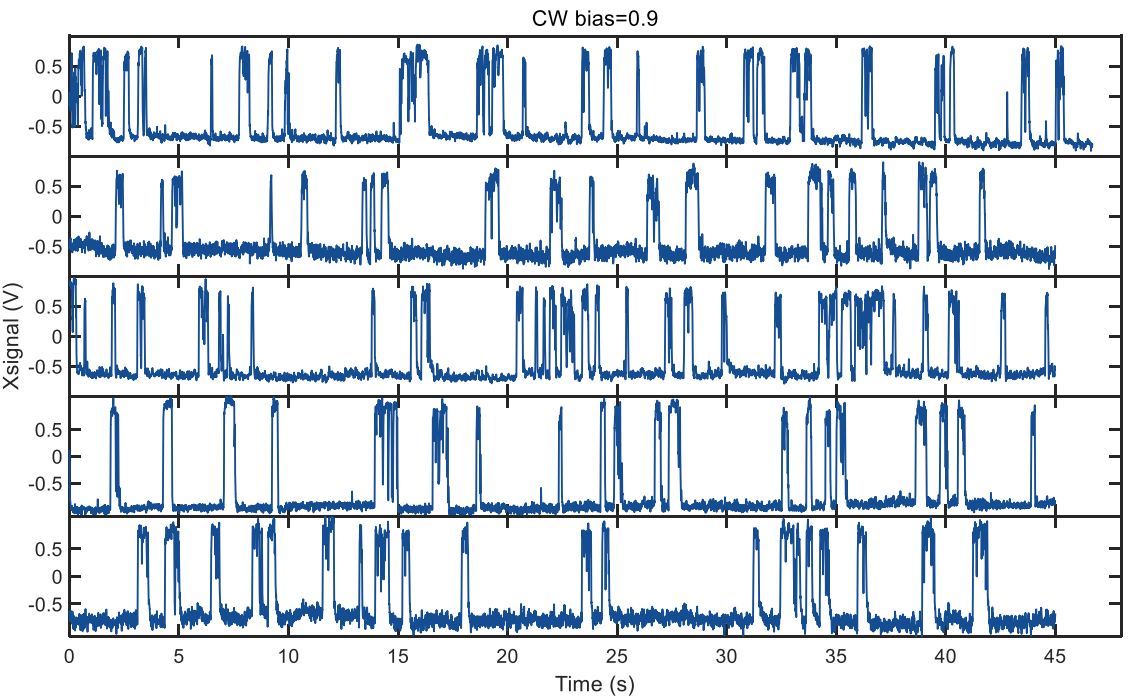

Fig. S1. More examples of motor switching traces at stall at various CW biases.

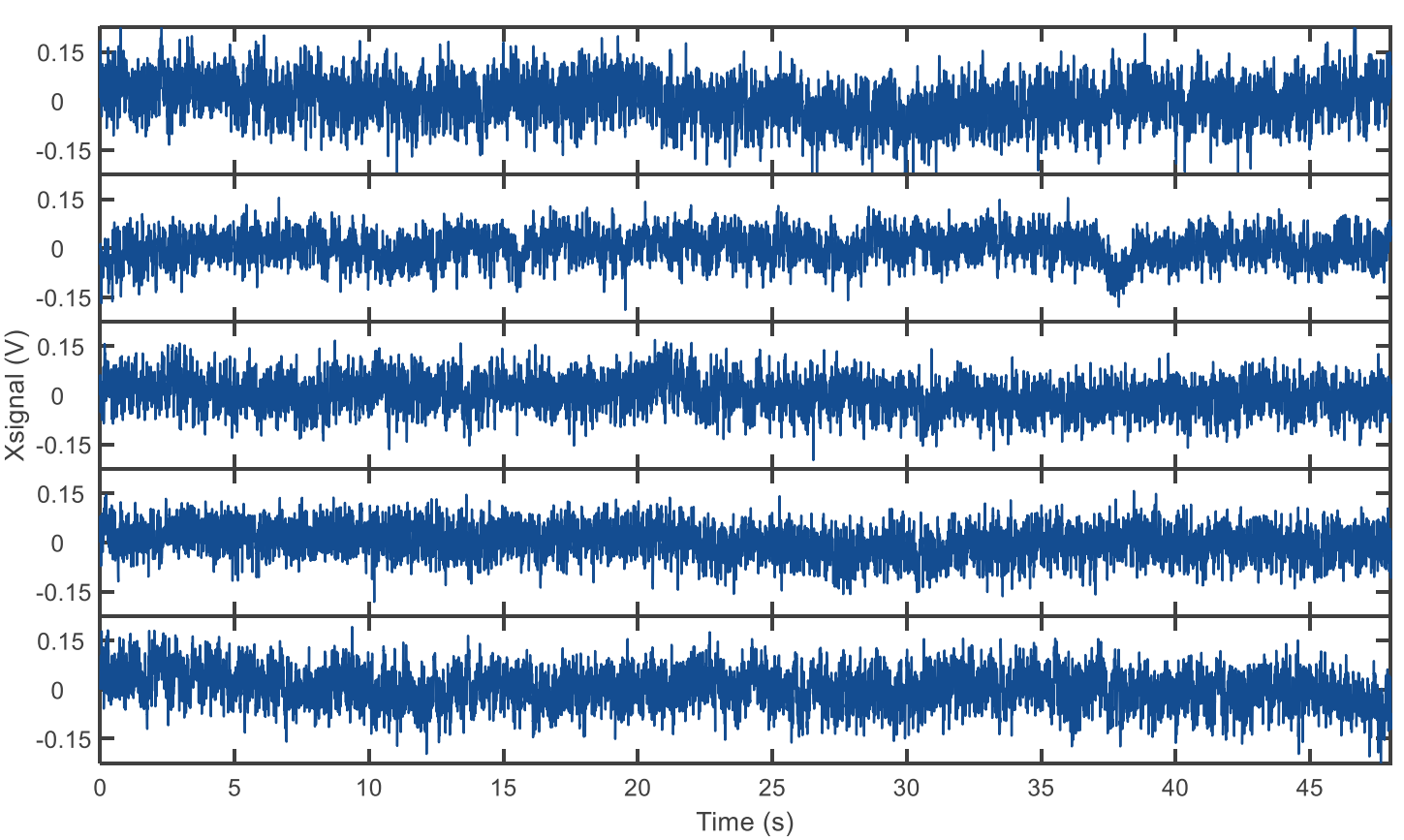

Fig. S2. The traces for stalled motors of the CCW-only strain JY9.

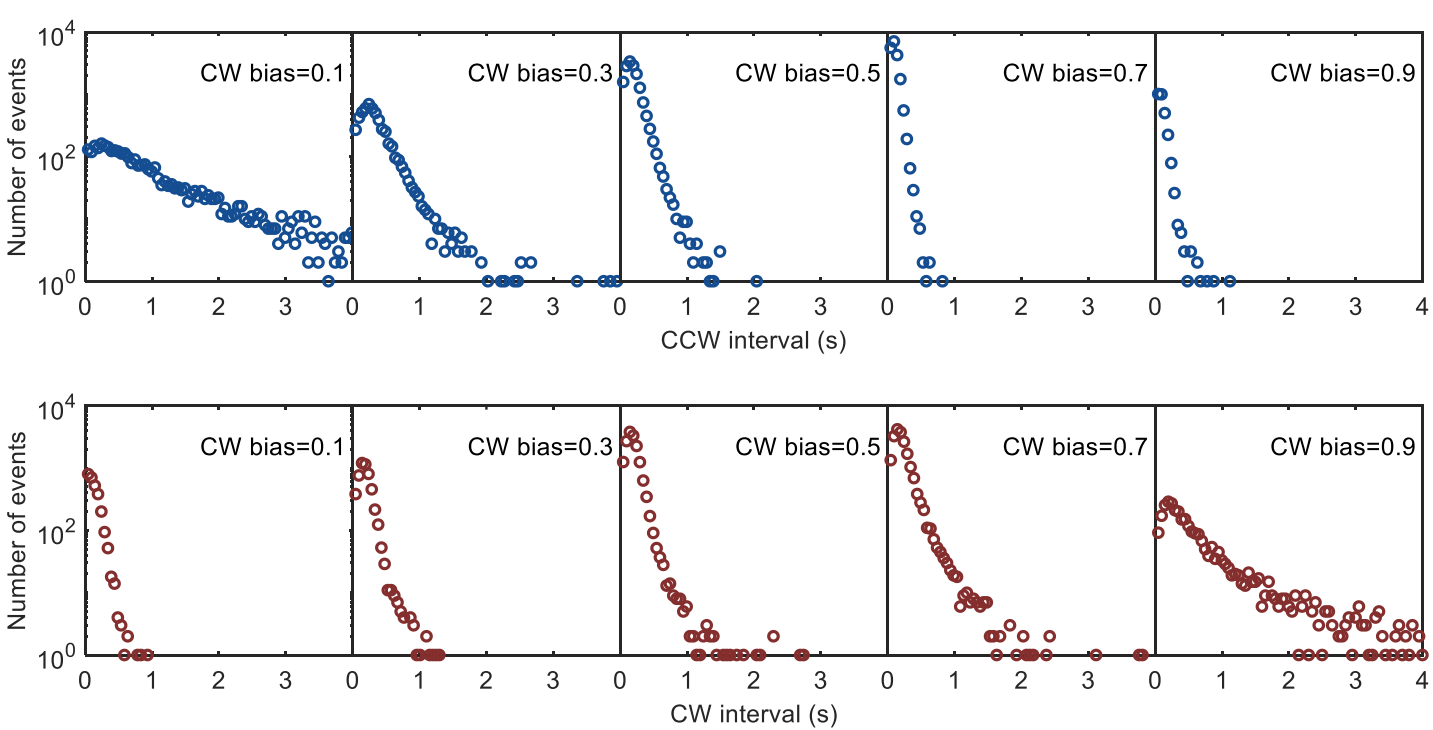
Fig. S3. CCW and CW interval distributions for motors driving the rotation of 750-nm-diameter bead on shortened filament stubs. 111 motors were observed, each for 200 s. The numbers of motors observed were 20, 14, 31, 32, and 14 for the groups centering on CW biases of 0.1, 0.3, 0.5, 0.7, and 0.9, respectively.

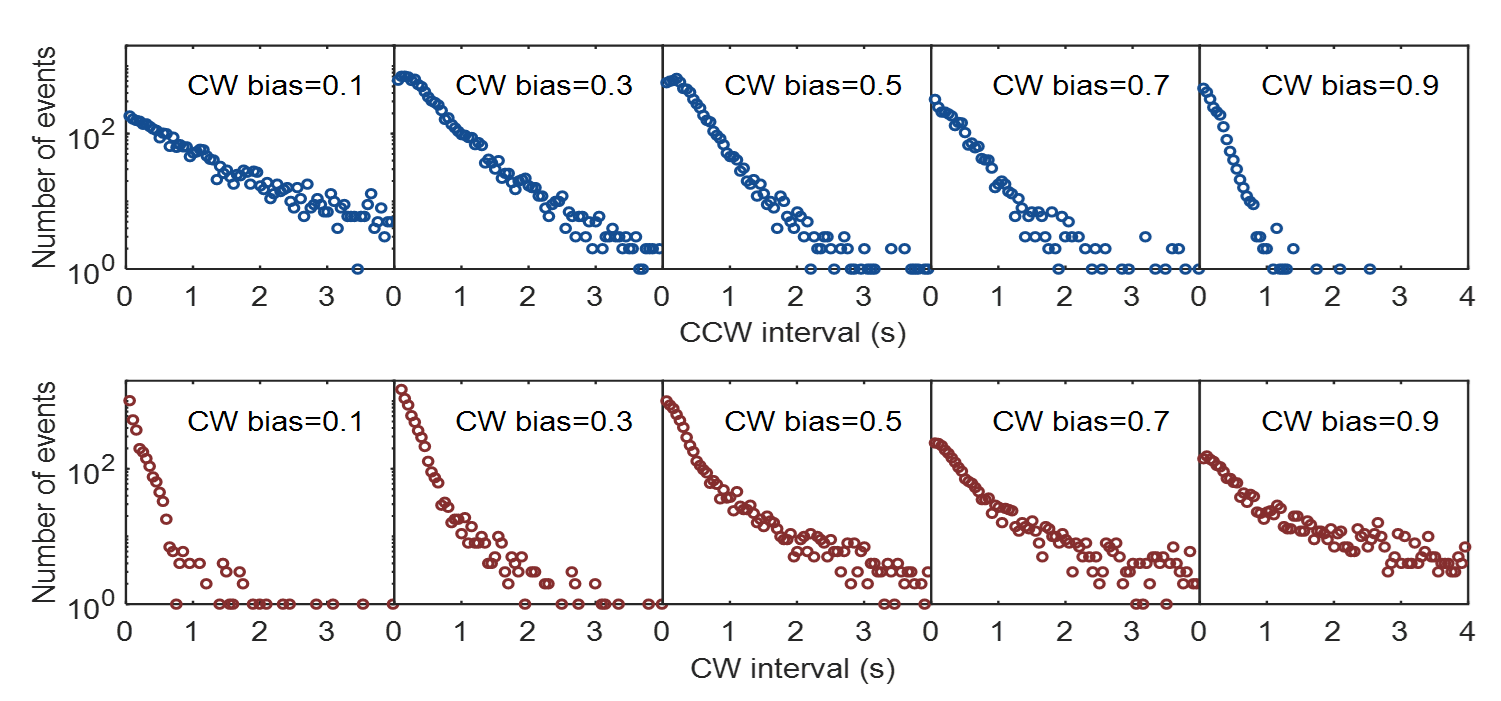

Fig. S4. CCW and CW interval distributions for motors driving the rotation of 1500-nm-diameter bead on shortened filament stubs. 135 motors were observed, each for 200 s. The numbers of motors observed were 26, 34, 28, 18, and 29 for the groups centering on CW biases of 0.1, 0.3, 0.5, 0.7, and 0.9, respectively.

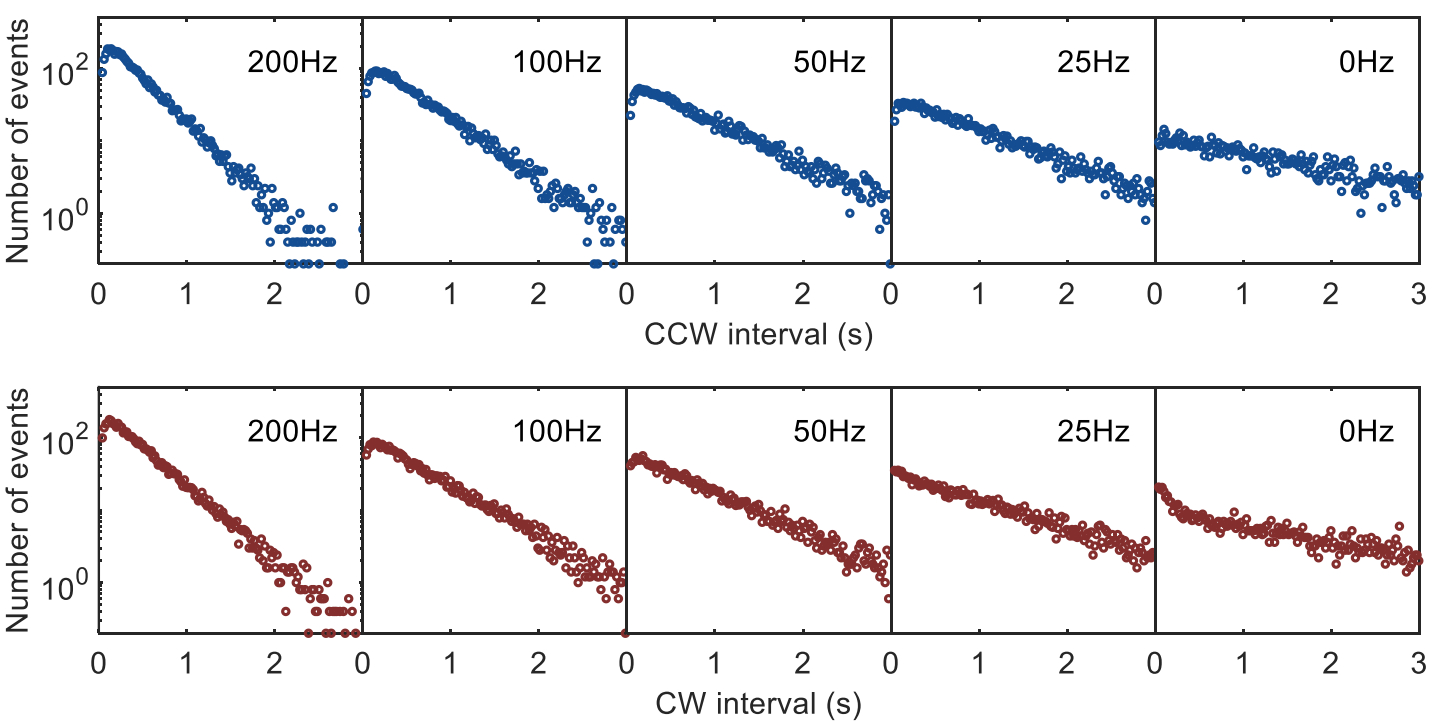

Fig. S5. The CCW (top) and CW (bottom) interval distributions for motor switching under different loads (at different CCW speeds) from our simulation with a CW bias of 0.5 and using *δ*_B_ = −*δ*_U_ =0.16. As the motor speed decreases from 200 Hz to stall, the shape of the distributions changes from non-exponential with a peak at short intervals to exponential.

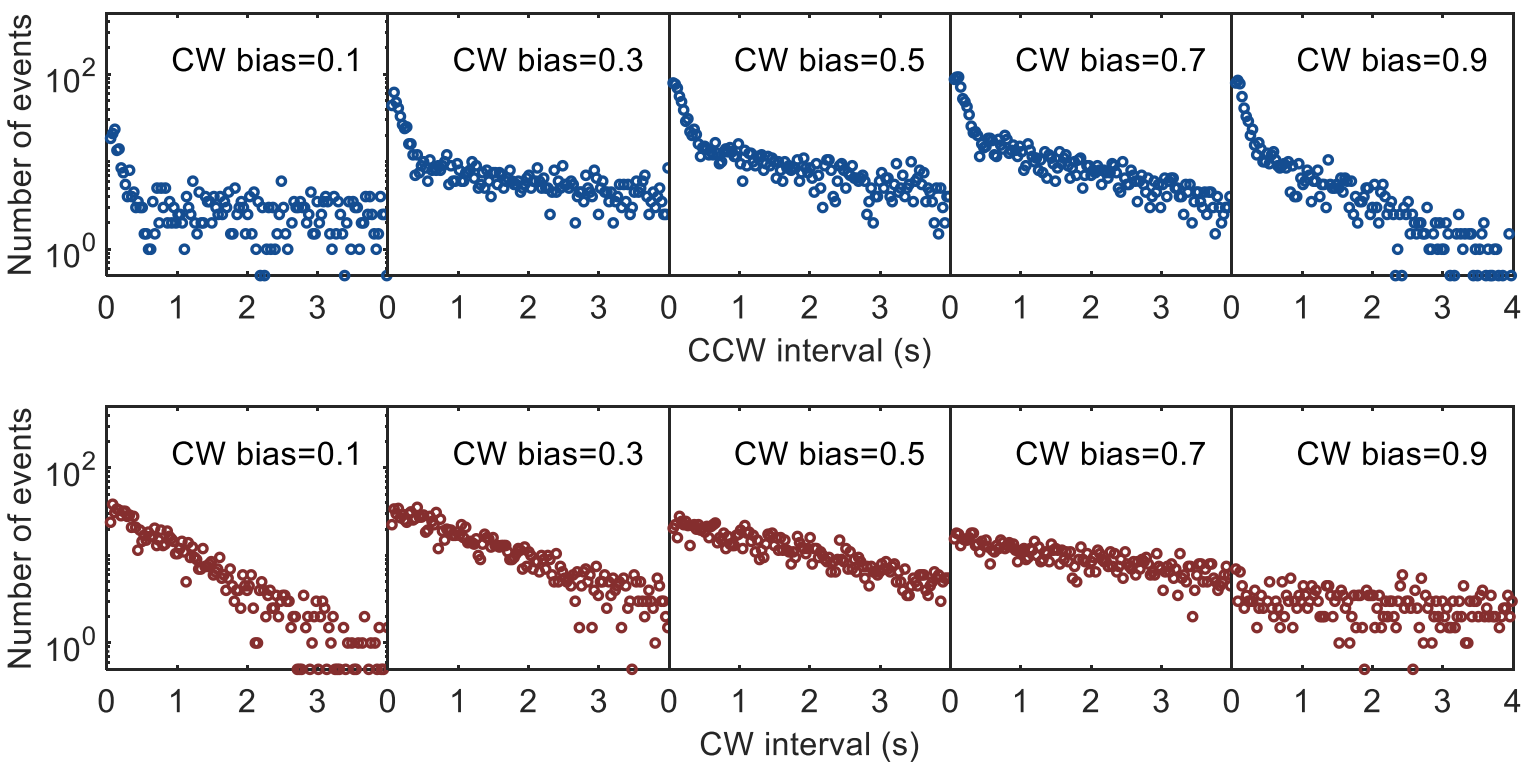

Fig. S6. The interval distributions at different CW biases for motor switching at stall from the simulation, using *δ*_U_ = −*δ*_B_ =0.16.
